# CD117/c-kit Defines a Prostate CSC-Like Subpopulation Driving Progression and TKI Resistance

**DOI:** 10.1101/256107

**Authors:** Koran S. Harris, Lihong Shi, Brittni M. Foster, Mary E. Mobley, Phyllis L. Elliott, Conner J. Song, Kounosuke Watabe, Carl D. Langefeld, Bethany A. Kerr

**Affiliations:** Department of Cancer Biology, Wake Forest School of Medicine, Winston-Salem, NC 27157; Wake Forest Baptist Comprehensive Cancer Center, Winston-Salem, NC 27157; Department of Biostatistics and Data Science, Division of Public Health Sciences, Wake Forest School of Medicine, Winston-Salem, NC 27157; Department of Urology, Wake Forest School of Medicine, Winston-Salem, NC 27157

**Author notes:** Corresponding Author: Bethany Kerr, Ph.D., Wake Forest School of Medicine, Medical Center Blvd, Winston-Salem, NC, 27157.; Twitter: @BethanyKerrLab. Abbreviations: CSC: cancer stem-like cell; CTC: circulating tumor cells; EMT: epithelial-mesenchymal transition; HSC: hematopoietic stem cell; SCF: stem cell factor; TKI: tyrosine kinase inhibitor.

**Keywords:** cancer stem cell, CD117/c-kit, SCF, migration, therapeutic resistance, circulating tumor cell, tyrosine kinase inhibitor

## Abstract

Cancer stem-like cells (CSCs) are associated with cancer progression, metastasis, and recurrence, and may also represent a subset of circulating tumor cells (CTCs). In our prior study, CTCs in advanced prostate cancer patients were found to express CD117/c-kit in a liquid biopsy. Whether CD117 expression played an active or passive role in the aggressiveness and migration of these CTCs remained an open question. In this study, we show that CD117 expression in prostate cancer patients is associated with decreased overall and progression-free survival and that activation and phosphorylation of CD117 increases in prostate cancer patients with higher Gleason grades. To determine how CD117 expression and activation by its ligand stem cell factor (SCF, kit ligand, steel factor) alter prostate cancer aggressiveness, we used LNCaP-C4-2 and PC3-mm human prostate cancer cells, which contain a CD117+ subpopulation. We demonstrate that CD117+ cells display increased proliferation and migration. In prostaspheres, CD117 expression enhances sphere formation. In both 2D and 3D cultures, stemness marker gene expression is higher in CD117+ cells. Using xenograft limiting dilution assays and serial tumor initiation assays, we show that CD117+ cells represent a CSC population. Combined, these data indicate that CD117 expression potentially promotes tumor initiation and metastasis. Further, in cell lines, CD117 activation by SCF promotes faster proliferation and invasiveness, while blocking CD117 activation with tyrosine kinase inhibitors (TKIs) decreased progression in a context-dependent manner. We demonstrate that CD117 expression and activation drives prostate cancer aggressiveness through the CSC phenotype and TKI resistance.

## INTRODUCTION

Prostate cancer is the second leading cause of cancer mortality in American men, with an estimated 191,930 new cases in 2020. When diagnosed locally and treated, patients can expect a 99% 5-year survival rate. Once cancer has metastasized or recurred; however, the 5-year survival rate drops to 30% driving ongoing research to identify and treat patients harboring aggressive tumors.[1] Currently, available nomograms are unable to distinguish patients with indolent disease from those likely to experience metastasis. Ongoing research has focused on understanding the individual steps of metastasis in an attempt to develop new methods to identify patients with aggressive disease. The first steps of the metastatic process: epithelial-mesenchymal transition (EMT), local invasion, intravasation, and survival in the circulation, represent early targets for identifying patients with aggressive tumors.[2]–[4] Cancer cells completing these steps become circulating tumor cells.

Circulating tumor cells (CTCs) are released at a rate of 3.2 x 10^6^ cells/g tissue daily, comprising one cell out of 10^5^ to 10^7^ leukocytes in the bloodstream.[3],[5],[6] However, <0.01% of these cells will initiate metastases.[3],[5] While multiple markers were proposed for prostate cancer CTCs,[7] most have yet to be validated in the circulation of patients. Further, it remains unproven whether CTCs can initiate metastases, which is partially due to an inability to examine the journey of CTCs from the primary tumor to the metastatic niche.[8] A subset of CTCs postulated to drive metastasis are cancer stem-like cells (CSCs).[9],[10] The CSC theory postulates that a subpopulation of tumor cells remaining after resection drives recurrence, while CTCs surviving the circulation and arresting at metastatic sites driving tumor growth are metastatic CSC.[11] In either case, CSCs are capable of self-renewal and asymmetric division and may be able to recapitulate the initial tumor heterogeneity. Further, these CSCs are more resistant to most treatments.[11]–[18]

Several markers for CTCs and CSCs are postulated in the literature.[7] In a prior study, we identified CD117/c-kit positive CTCs in prostate cancer patients that were associated with cancer severity and biochemical recurrence.[19] CD117+ cell numbers were increased in the prostate tissue and circulation of patients with Gleason 8+ disease. Further, three months postoperatively, CD117 CTC numbers were equal or increased in patients with recurrent cancer, while CD117 + cell numbers had dropped 50% in patients for whom surgery was successful.[19] Further studies demonstrated that CD117 expression in prostate cancers was increased with Gleason score and that CD117 was expressed by both stromal cells in the transitional zone and cancer cells in the peripheral zone.[20] Additionally, CD117 is expressed on both normal stem cells and aggressive cancer cells with CD117 expression found on hematopoietic stem cells (HSCs) and stem cells in the murine prostate.[21]–[23] A single CD117 positive cell, which was also Lin-Sca-1+ CD133+ CD44+, regenerated an entire secreting prostate when mixed with urogenital mesenchymal cells and implanted in the renal capsule. Thus, this CD117 expressing cell was considered a prostate stem cell in adult tissue.[22] Thus, CD117 expression is associated with stemness in multiple cell types.

CD117 is a member of the type III tyrosine kinase receptors which play a key role in cell signaling and are responsible for maintaining cell functions such as cell survival, metabolism, cell growth and progression, proliferation, apoptosis, cell migration, and cell differentiation.[21],[24]–[26] These effects of CD117 activation and signaling support its potential role in prostate cancer progression and CSC maintenance. How CD117 activation alters proliferation, stemness, and migration remains to be elucidated. The present study aimed to determine the extent to which the CTC marker CD117 was involved in prostate cancer progression, CSC maintenance, and therapeutic resistance. The human prostate cancer cell lines LNCaP-C4-2 and PC3-mm were separated into CD117+ and negative populations and the differences between the two subpopulations examined. The effects of CD117 activation by SCF treatment or inhibition by tyrosine kinase inhibitor (TKI) treatment were also tested. Using live-cell imaging and xenograft models, we determine that CD117 activation drives progression, stemness, migration, and TKI resistance.

## METHODS

### Genomic Databases

RNA sequencing data from 4,638 individuals from the prostate cancer cohort were obtained from The Cancer Genome Atlas via cBioPortal (https://www.cbioportal.org/).[27],[28] Survival data was calculated by cBioPortal. Affymetrix mRNA expression, copy number, and protein expression data from prostate cancer cell lines were obtained from the Broad Institute Cancer Cell Line Encyclopedia (https://portals.broadinstitute.org/ccle).[29],[30]

### Flow Cytometry

To assess, the numbers of CD117 expressing cells in human prostate cancer cell lines, LNCaP-C4-2 (RRID: CVCL_4782), PC-3 (RRID: CVCL_0035), DU145 (RRID: CVCL_0105) (the prior three lines were gifts from Dr. Warren D. Heston, Cleveland Clinic), LNCaP-C4-2B (RRID: CVCL_4784; a gift from Dr. Magda Grabowska, Case Western Reserve University, via Dr. Simon Hayward and Dr. Leland Chung) and PC3-mm (RRID: CVCL_4885) were blocked with human FcR blocking reagent (1:100; Miltenyi Biotec) and subsequently incubated with anti-CD117/c-kit-APC (1:50; Miltenyi Biotec; RRID: AB_615056). Cells were fixed in 1% formalin and analyzed on a BD FACS Canto II running the FACS Diva Software (BD Biosciences). Fluorescence values were normalized to an unstained control samples and initial compensation applied using IgG-APC control antibody (Miltenyi Biotech; RRID: AB_871704).

### Cell Culture, Sorting, and Treatment

To examine how CD117 expression alters cell function, human prostate cancer LNCaP-C4-2 or PC3-mm cells were sorted into CD117+ and negative populations. LNCaP-C4-2 and PC3-mm cells were cultured in RPMI1640+10%FBS and confirmed negative for mycoplasma every six months. LNCaP-C4-2 cells were transduced to express ZsGreen or Firefly luciferase-mCherry by the Wake Forest Baptist Comprehensive Cancer Center (WFBCCC) Cell Engineering Shared Resource using a lentivirus (Clontech or Genecopoeia). LNCaP-C4-2 and PC3-mm were sorted into CD117+ and negative populations using the Miltenyi MACS sorting beads or the ThermoFisher MagniSort CD117 (c-kit) positive selection kit. Sorted cells were treated with 50 ng/mL recombinant stem cell factor (SCF; Miltenyi Biotec), 5 μM imatinib mesylate (Sigma), 5 μM sunitinib malate (Sigma), or 5 μM ISCK03 (Sigma).

### IncuCyte Live Cell Imaging

To image cells over time, an Essen Bioscience IncuCyte ZOOM live-cell imager was used for proliferation, scratch migration, chemotaxis, and sphere formation assays. For proliferation assays, sorted cells were plated at 2,000 cells per well and imaged over three days. The percentage of confluence was calculated. For scratch migration assays, sorted cells were plated at 50,000 cells per well. The next day, cells were scratched using the Essen Biosciences WoundMaker tool. Scratches were imaged for 48 hours and wound width and percent closure were measured. For chemotaxis assays, sorted cells were plated at 1,000 cells/top of the insert and imaged for 48 hours. The number of cells on the top and bottom of the insert were counted. For sphere formation assays, 1,000 sorted cells were plated in ultra-low attachment plates and imaged over nine days. The sphere area was calculated over time. Separate groups were treated with inhibitors or starved overnight before SCF treatment.

### Immunofluorescence

For detection of stem cell and EMT marker expression, immunofluorescent staining for; SOX2, OCT4, E-cadherin, N-Cadherin, and vimentin was conducted. Spheres were embedded into OCT freezing medium and sectioned at 10 μm thick onto UltraClear Plus Microslides (Denville Scientific). Sections were then fixed in 4% paraformaldehyde (PFA) and then incubated with either; goat polyclonal anti-SOX2 (1:20; R&D Systems; RRID: AB_355110), rabbit polyclonal anti-OCT4 (1:100; Abcam; RRID: AB_445175), goat polyclonal anti-E-cadherin (1:100; R&D Systems; RRID: AB_355568), rabbit polyclonal anti-N-cadherin (1:250; Abcam; RRID: AB_444317), or rabbit monoclonal anti-Vimentin (1:500; Abcam; RRID: AB_10562134). Sections were then incubated with either Alexa Fluor 555 polyclonal donkey anti-goat (1:500; Invitrogen; RRID: AB_2535853) or Alexa Fluor 555 polyclonal goat anti-rabbit secondary antibodies, respectively (1:500; Invitrogen; RRID: AB_2535850). Slides were mounted with Prolong Gold Antifade DAPI (1:5000; Invitrogen). Images were captured on an Olympus VS110 at 10x by the Virtual Microscopy Core, and relative fluorescent was quantified using ImageJ software (v1.80; NIH).

### Quantitative PCR Gene Expression

To examine gene expression, cells and spheres were lysed using the Qiagen RNeasy kit and cDNA synthesized using the Applied Biosystems High Capacity cDNA Reverse Transcription kit. Primer sequences are listed in Table 1. For detection and quantification, PCR reactions were run on a Roche Light Cycler 480II using the Applied Biosystems SYBR Green PCR Master Mix. Data were analyzed by the ΔΔCT method.

**Table 1.**
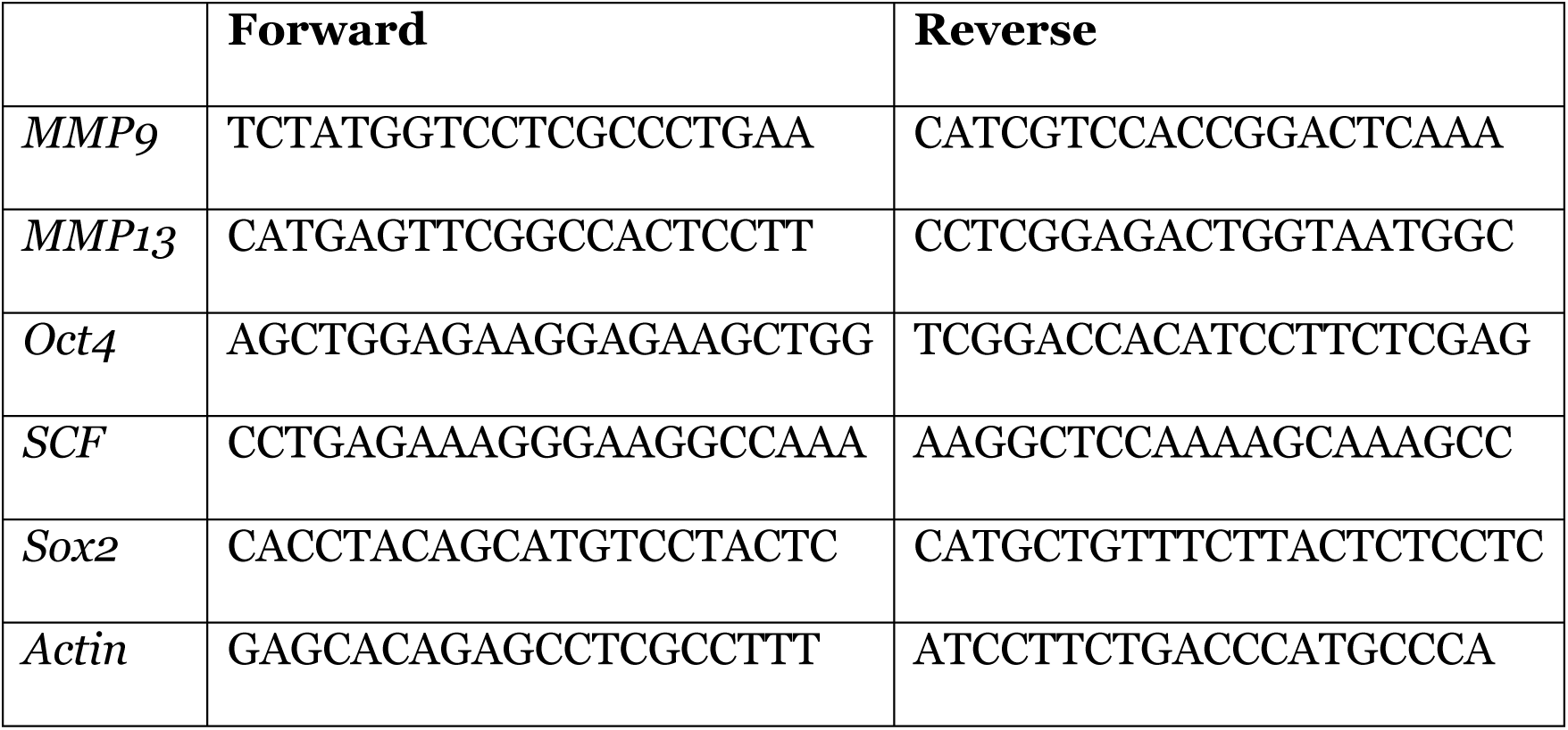
qPCR Primers. Primer sequences for genes examined by quantitative PCR.

### Tumor Initiation

To assess tumor-initiating capabilities, NOD.CB17-Prkdc^scid^ mice (Jackson Laboratory; RRID: IMSR_JAX:001303) aged between 8 and 12 weeks old were injected subcutaneously with decreasing numbers of sorted LNCaP-C4-2 cells (10,000 to 10 cells) under Wake Forest School of Medicine IACUC Approval #A15-221. Tumors were imaged weekly using an IVIS Lumina bioluminescent imager after injection with 150 mg/kg luciferin (WFBCCC Cell Engineering Shared Resource). After 30 days, tumors were removed and weighed. For serial tumor initiation, tumors were dissociated using the Miltenyi tumor dissociation kit run on the Miltenyi gentleMACS Octo Dissociator. Resulting cells were again bead isolated and injected into a second group of mice at 10 cells and allowed to grow 30 days before tumor removal.

### Tumor Microarray Analysis

To assess whether CD117 was activated in prostate cancers, we obtained tissue microarrays (TMAs) from the WFBCCC Tumor Tissue and Pathology Shared Resource under IRB Approval #38212. TMAs processed from immunohistochemistry to visualize phosphorylated (p-) CD117 (R&D Systems, 1:50, RRID: AB_1152039) and prostate-specific membrane antigen (PSMA) (Agilent, 1:50, RRID: AB_2106450) in tissues. Slides were scanned on a Hamamatsu Photonics NanoZoomer by the Virtual Microscopy Core. Staining was quantified using Indica Labs HALO software and grouped according to Gleason Score.

### Statistical Analysis

To determine statistical significance, Student’s *t* test or one-way analysis of variance with Tukey post-test were used to analyze data with the GraphPad Prism 7 software (RRID: SCR_002798). For experiments over time, linear regression models were developed to test for differences in slopes over time and test interactions between cell type and slope of the relationship with time using the SAS software. Error bars represent the SEM of experiments. * p<0.05, ** p<0.01, and *** p<0.005. Experiments were repeated at least three times.

## RESULTS

### CD117 Expression Results in Decreased Survival

In a prior study, we demonstrated that the number of CD117+ cells circulating in prostate cancer patients was associated with cancer severity and biochemical recurrence.[19] Based on these findings, we asked how CD117 expression on these cells promoted prostate cancer progression. In an initial study, we showed that CD117+ LNCaP-C4-2 xenograft tumors were larger and more angiogenic than tumors containing the negative population.[19] To examine the outcomes of prostate cancer patients bearing CD117 alterations in their tumors, we used the cBioPortal database to mine the TCGA database. CD117 expression was found to be amplified in patients in 11 of the 19 studies examined representing 0.6% to 16.2% of the patients. Median overall survival time was reduced to 77 months in patients with CD117 alteration compared with 133 months (Logrank Test p = 5.23 x 10^-7^) for unaltered patients (Figure 1A). Additionally, progression-free survival was decreased in patients with CD117 gene alterations (Logrank Test p = 8.951 x 10^-3^; Figure 1B). We then used the Broad Institute Cell Line Encyclopedia to examine changes in CD117 expression in prostate cancer cell lines. The mRNA for CD117 was highly expressed in all cell lines with lower protein expression (Figure 1C). Accordingly, we profiled several prostate cancer cell lines including several bone-trophic lines not currently available in the Cell Line Encyclopedia for CD117 cell surface expression by flow cytometry. We discovered a small CD117+ population in PC-3 (∼0.61%) and DU145 (∼0.685%) prostate cancer cell lines. The bone-trophic PC3-mm cell line contained a ∼2% CD117+ subpopulation. The androgen-independent and bone-trophic LNCaP-C4-2 and LNCaP-C4-2B prostate cancer cell lines (derived from LNCaP cells) both contained significant CD117+ populations, ∼21.4% and ∼32.3% on average respectively.

**Figure 1.**
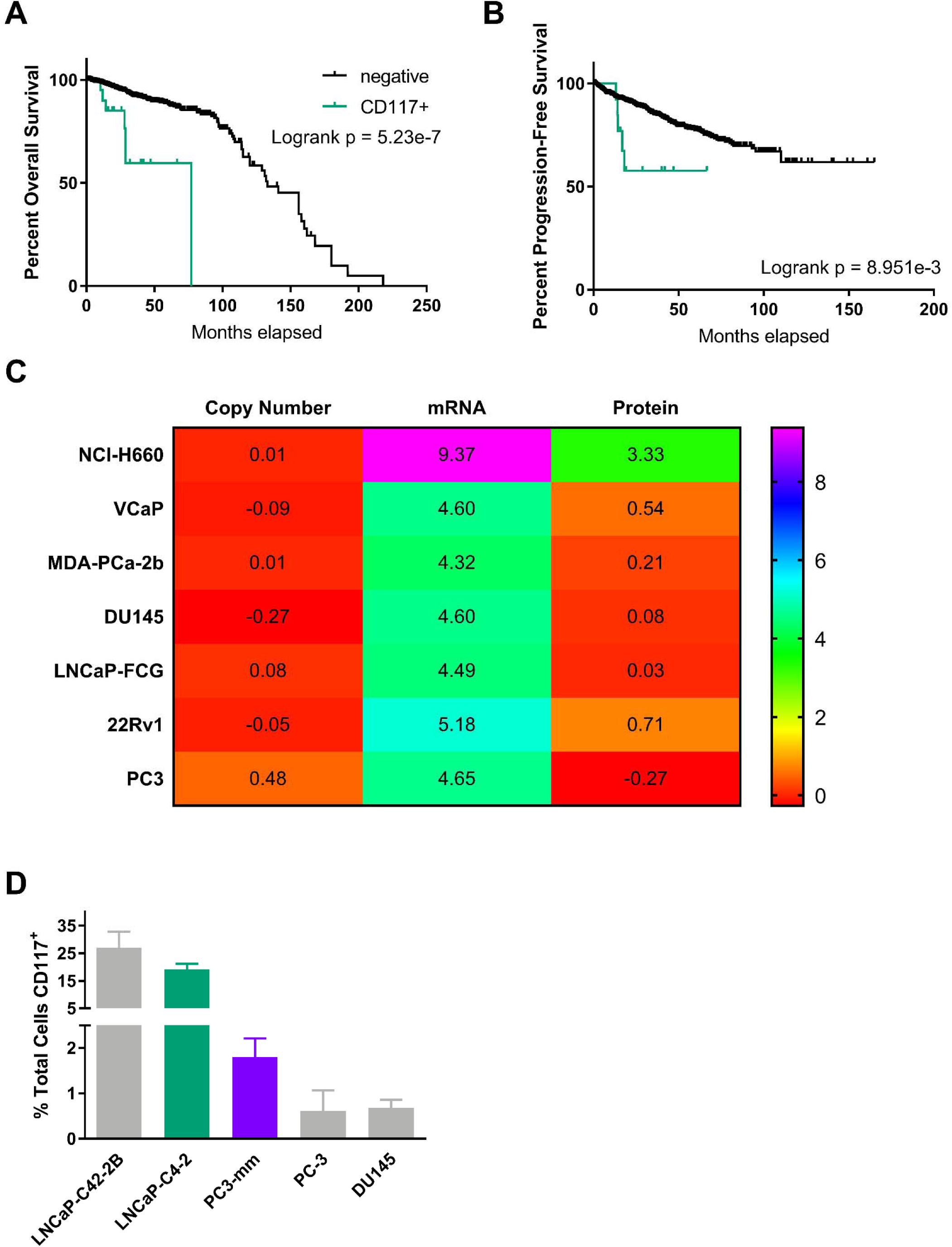
CD117 Expression is Associated with Worse Survival and is Found in Prostate Cancer Cell Lines. (A-B) Overall survival (*n*=1,476 patients) and progression-free survival (*n*=1,329 patients) percentages with CD117 alteration were obtained from The Cancer Genome Atlas via cBioPortal. (C) Copy number, mRNA expression, and protein expression values for CD117 in several prostate cancer cell lines were obtained from the Broad Institute Cancer Cell Line Encyclopedia. (D) The percentages of CD117+ cells were measured by flow cytometry for LNCaP-C4-2B, LNCaP-C4-2, PC3-mm, PC-3, and DU145 prostate cancer cell lines and represented as mean ±SEM (*n*=3).

### CD117 Expression Stimulates Prostate Cancer Proliferation and Migration

To further determine how CD117 expression drives prostate cancer aggressiveness, we sorted LNCaP-C4-2 and PC3-mm human prostate cancer cells into CD117+ and negative populations. Using live-cell imaging, we examined differences between these two populations. In a proliferation assay, CD117+ cells proliferated more quickly than the negative cells. LNCaP-C4-2 CD117+ cells reached confluency peak at hour 58 at which point CD1117+ cells were 2.9-fold more confluent than the negative population (Figure 2A). Similarly, PC3-mm CD117+ cells at hour 58 at were 1.75-fold more confluent than the negative population (Figure 2B) Thus, in monolayer culture, CD117 promotes proliferation. Because the LNCaP-C4-2 prostate cancer cell line contained a higher CD117+ subpopulation, we profiled them in the rest of our studies.

**Figure 2.**
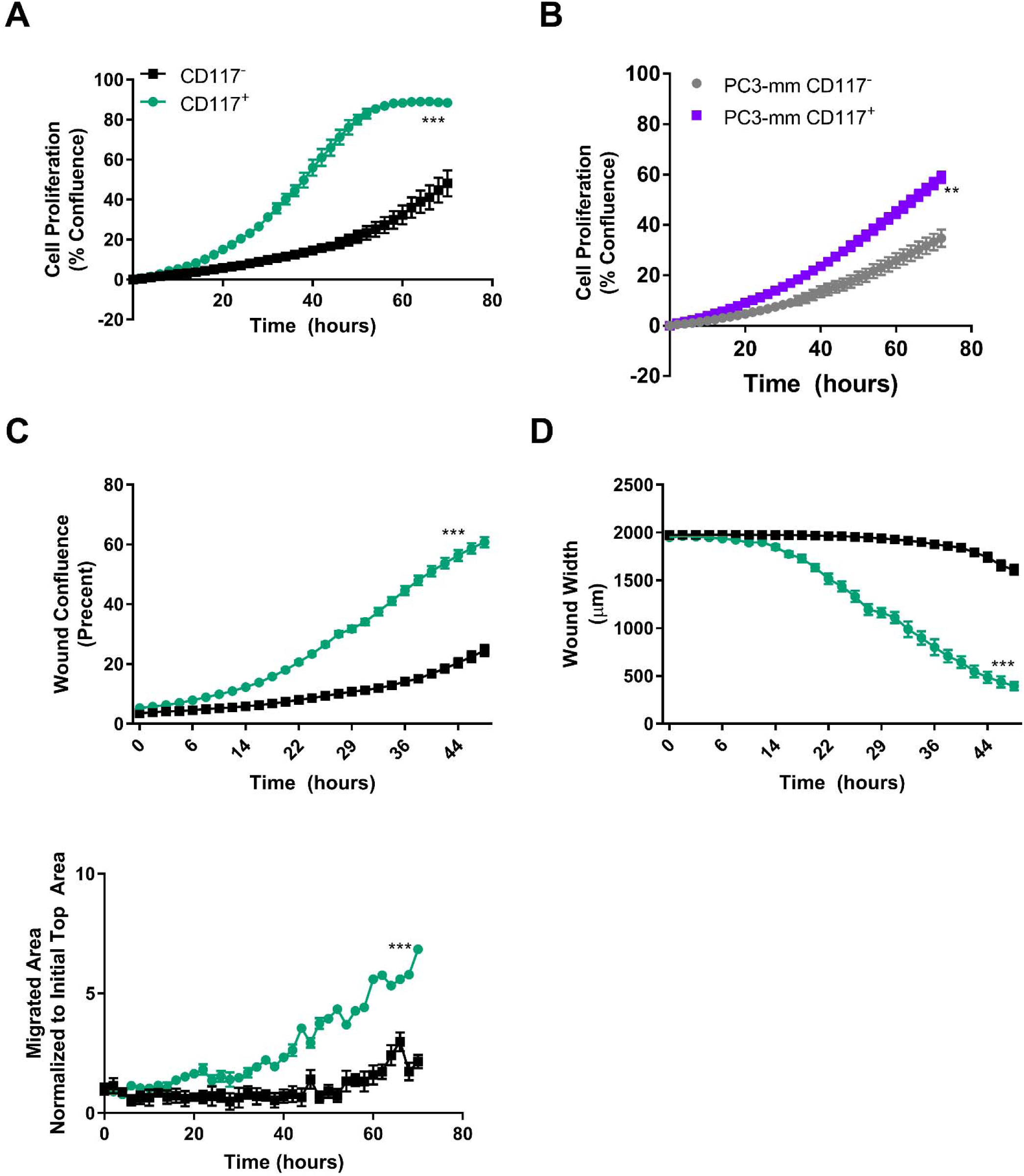
CD117 Expression Induces Cancer Cell Proliferation and Migration. (A) LNCaP-C4-2 cells were sorted into CD117+ and negative populations, proliferation measured by live-cell imaging, and represented as mean percent confluence ±SEM (*n*=12). (B) PC3-mm cells were sorted into CD117+ and negative populations, proliferation measured by live-cell imaging, and represented as mean percent confluence ±SEM (*n*=12). (C-D) LNCaP-C4-2 cells were sorted into CD117+ and negative populations. Cell migration was measured by live-cell imaging of a scratch assay. Percent cell confluence in the wound (C) and wound width (D) are represented as mean ±SEM (*n*=6). (E) Chemotaxis through pores as measured by live-cell imaging and represented as number of cells on transwell bottom normalized to initial seeding mean ±SEM (*n*=8). *** represents p<0.005 by Student’s *t* test.

Since CD117+ cells were found in the patient circulation, we examined cancer cell migration using scratch and chemotaxis assays. Confluent LNCaP-C4-2 CD117+ and negative cells were scratched to generate a wound, and cell movement into the wound was measured by live-cell imaging. CD117+ cultures showed increased confluence within the wound and decreased wound width over time (Figure 2C and D). Upon experimental termination, CD117+ cell wound closure was 2.7-fold higher, while the wound width was 1.7-fold smaller compared to the negative population. Additionally, CD117+ and negative cells were plated on the top of a transwell, and migration through pores measured over time. At experimental termination, 3.2-fold more CD117+ cells had migrated when compared to the negative population (Figure 2E). Thus, CD117 expression increases cell migration in two dimensions.

Another marker of aggressive tumor cells is the ability to form spheres.[17],[31]–[35] Sorted cells were examined for sphere formation by live-cell imaging and changes in gene and protein expression of aggressiveness markers was measured in spheres. CD117+ cells formed 1.35-fold larger spheres on day 5 compared with the negative cells (Figure 3A). However, staining for the EMT marker vimentin demonstrated a 1.7-fold increase in CD117+ spheres compared with negative (Figure 3B). Additionally, a 2.0-fold increase in the stemness marker Oct4 was measured in CD117+ spheres (Figure 3B). To examine how growth in 3D spheres altered gene expression, we compared samples to 2D, monolayer grown cultures. The growth of CD117+ cells in spheres upregulated *Oct4* (6.8-fold) and *MMP-13* (7.7-fold) compared with monolayer cultures (Figure 3C). Interestingly, the ligand for CD117, *SCF*, expression was also increased with 3D growth of the CD117+ cells. (Figure 3C). No significant difference was seen between negative culture conditions (Figure 3C). Taken together, these data demonstrate that CD117 expression may promote prostate cancer progression and this effect is increased when cells are grown in 3D spheres.

**Figure 3.**
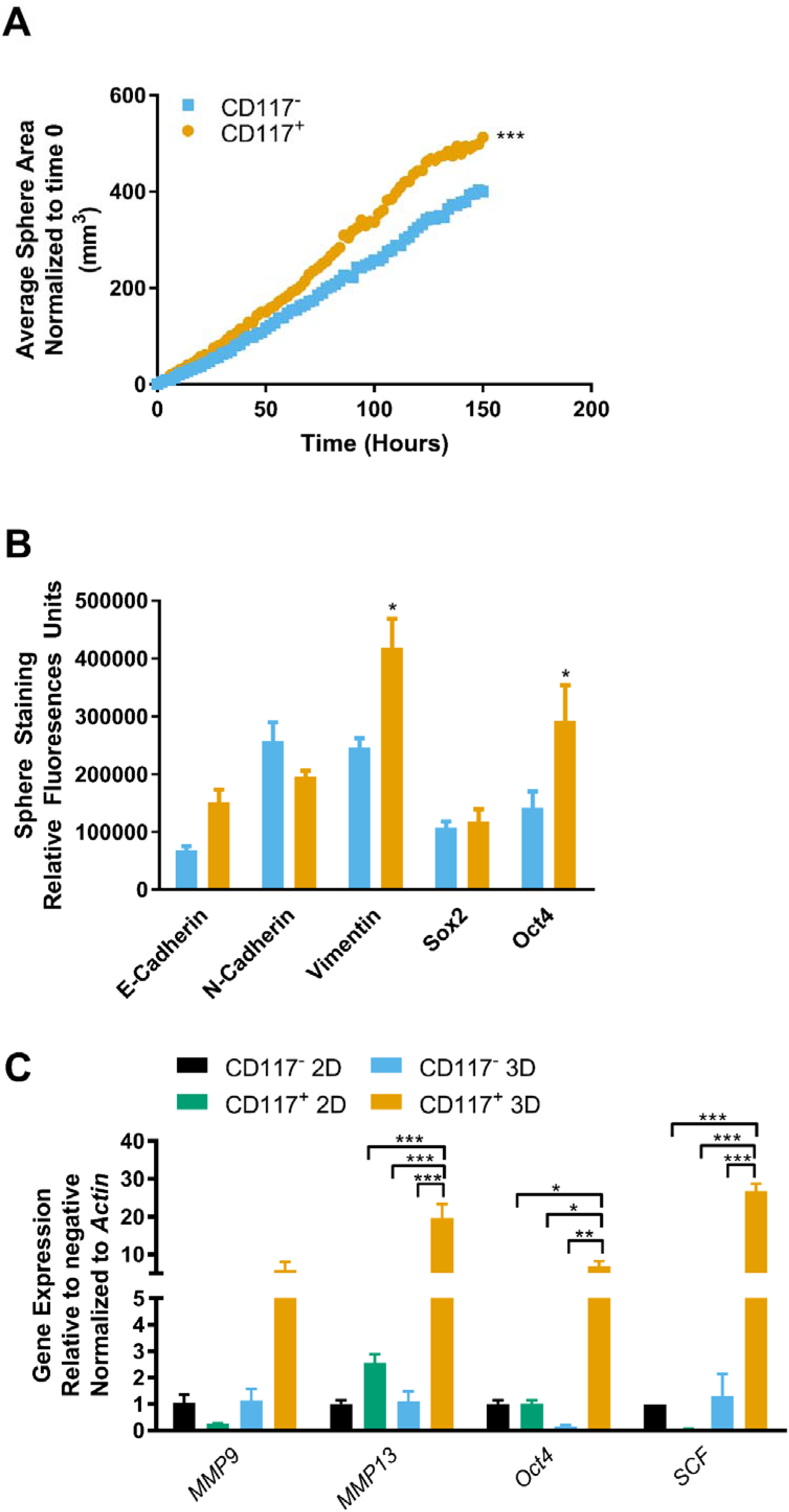
CD117 Promotes Aggressiveness in Three-Dimensional Culture. LNCaP-C4-2 cells were sorted into CD117+ and negative populations. (A) Sphere formation was tracked using live-cell imaging and represented as mean sphere area (*n*=12). (B) After 7 days, spheres were sectioned and stained for the EMT markers: E-cadherin, N-cadherin, and vimentin; and the stemness markers: Sox2 and Oct4. Relative fluorescence is represented as mean ±SEM (*n*=3-6). (C) Gene expression was compared between sorted cells grown in 2D monolayer or 3D spheres and represented as mean fold change ±SEM (*n*=3). * represents p<0.05, ** represents p<0.01, and *** represents p<0.005 by Student’s *t* test (A-B) and one-way ANOVA (C).

### CD117+ Cells Are Cancer Stem-like Cells

More invasive cancer cells that form prostaspheres have the potential to be CSCs.[17] Additionally, the expression of stem cell markers, including *Oct3/4* and *Sox2* are often used to examine stemness in potential CSCs.[31],[36],[37] To further demonstrate pluripotency, self-renewal capacity is measured by clonogenic assays and serial *in vivo* tumor initiation or limiting dilution experiments designed to examine whether a population could regenerate an entire tumor and thus be considered a CSC.[13],[14],[16],[38],[39] In order to differentiate tumor-initiating cells from CSCs, repeated tumor-initiating xenografts are required.[39],[40] To determine whether CD117+ cells displayed the CSC phenotype, we first examined the expression of stemness markers. Gene expression of *Oct4* and *Sox2* were 2.5- and 15.5-fold higher, respectively, in CD117+ cells compared with the negative population (Figure 4A). To validate if these cells were a true cancer stem-like population, we performed serial tumor initiation experiments. CD117+ and negative cells were implanted subcutaneously in immunocompromised mice at limiting dilutions. Both populations formed tumors from 500-10,000 implanted cells (data not shown). At 100 cells both groups reliably formed tumors, while at 25 cells approximately 40% of CD117+ and 30% of negative cell implants formed tumors (Figure 4B). At 10 cells, 50% (13 of 26) of CD117+ and 23% (6 of 26) of negative cell implantations formed tumors (Figure 4B). At all cell concentrations, CD117+ cells formed larger tumors (Figure 4C). Serial tumor initiation was completed by dissociating the 10 cell tumors, repeating bead sorting, and re-implantation at 10 cells. In serial tumor initiation, CD117+ cells formed a second tumor ∼25% of time (3 of 20 from 10 cells and 7 of 20 from 25 cells), while the negative implants were unable to initiate secondary tumors (Figure 4D). Thus, CD117 positive cells represent a prostate cancer stem-like subpopulation.

**Figure 4.**
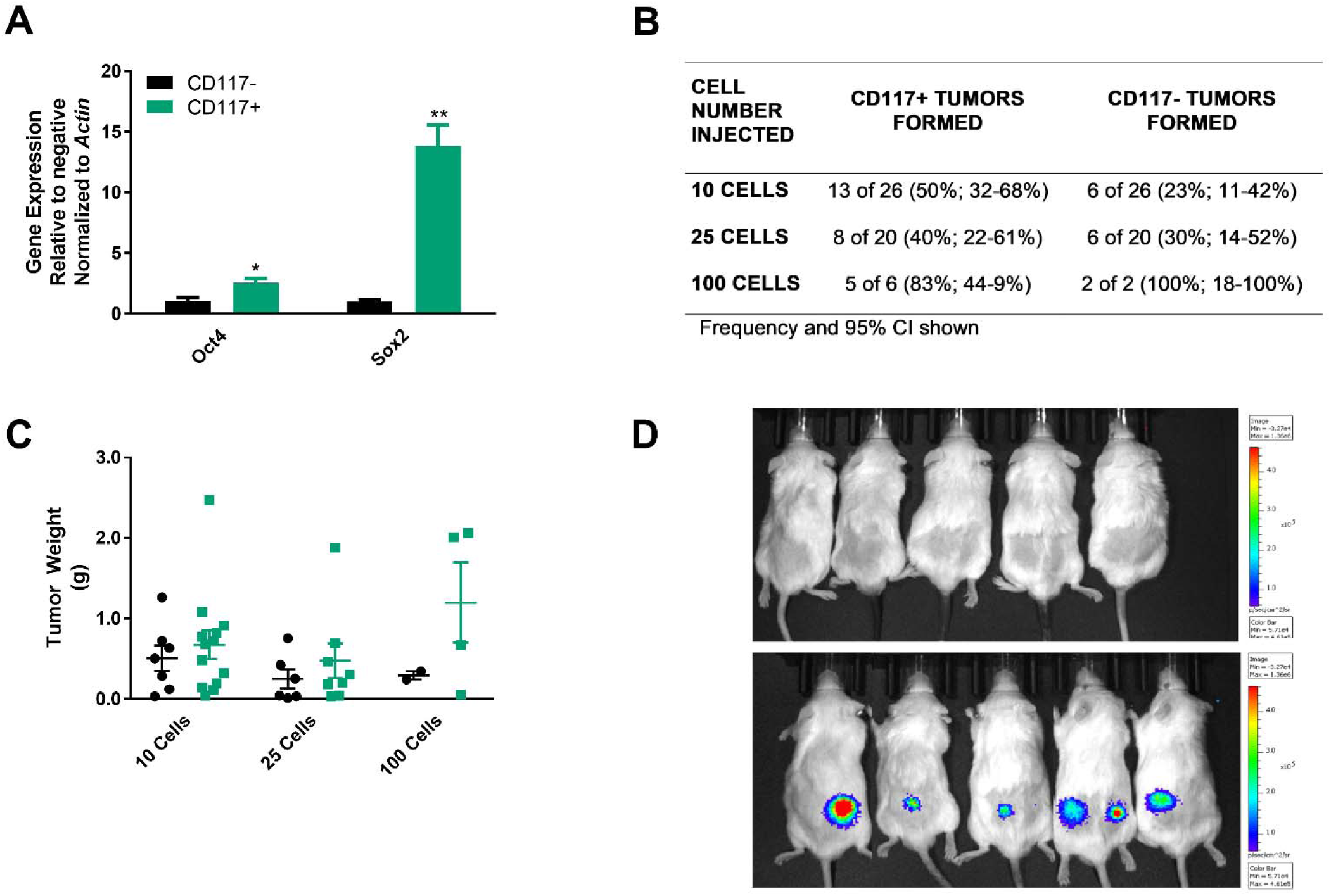
CD117+ Cells Represent a Cancer Stem-like Cell. LNCaP-C4-2 cells were sorted into CD117+ and negative populations. (A) Gene expression of monolayer cultured cells was examined for the stemness markers *Oct4* and *Sox2* and represented as mean fold change ±SEM (*n*=4). (B-D) Sorted cells were injected into immunocompromised mice in a limiting dilution assay. The number of tumors formed at each dilution is shown in (B) with the frequency percentage and 95% confidence intervals in parentheses. Tumor weights (C) are represented as mean ±SEM (*n*=2-13). IVIS imaging was performed to track tumor size and presence over time. A representative image is shown in (D). * represents p<0.05 and ** represents p<0.01 by Student’s *t* test.

### SCF Activation of CD117 Increases Prostate Cancer Progression

Based on these data demonstrating that CD117 expression drives prostate cancer progression, we questioned how activation of CD117 might alter these effects. CD117 is activated by binding to SCF, its sole ligand. SCF, found in a dimer, binds to CD117 inducing dimerization, phosphorylation, and downstream signaling leading to proliferation, cytoskeletal rearrangement, and migration.[21],[41],[42] To examine how activation of the tyrosine kinase receptor CD117 may alter prostate cancer aggressiveness, we first examined phosphorylated (p-)CD117 levels in prostate cancer patients. The numbers of p-CD117+ cells per core increased with Gleason Score (Figure 5A). Tumors from patients with grade 4 and 5 cancers contained 2.2-fold more p-CD117 cells compared with grade 1 patient tumors. Thus, activated CD117 is increased in patients with cancer severity.

**Figure 5.**
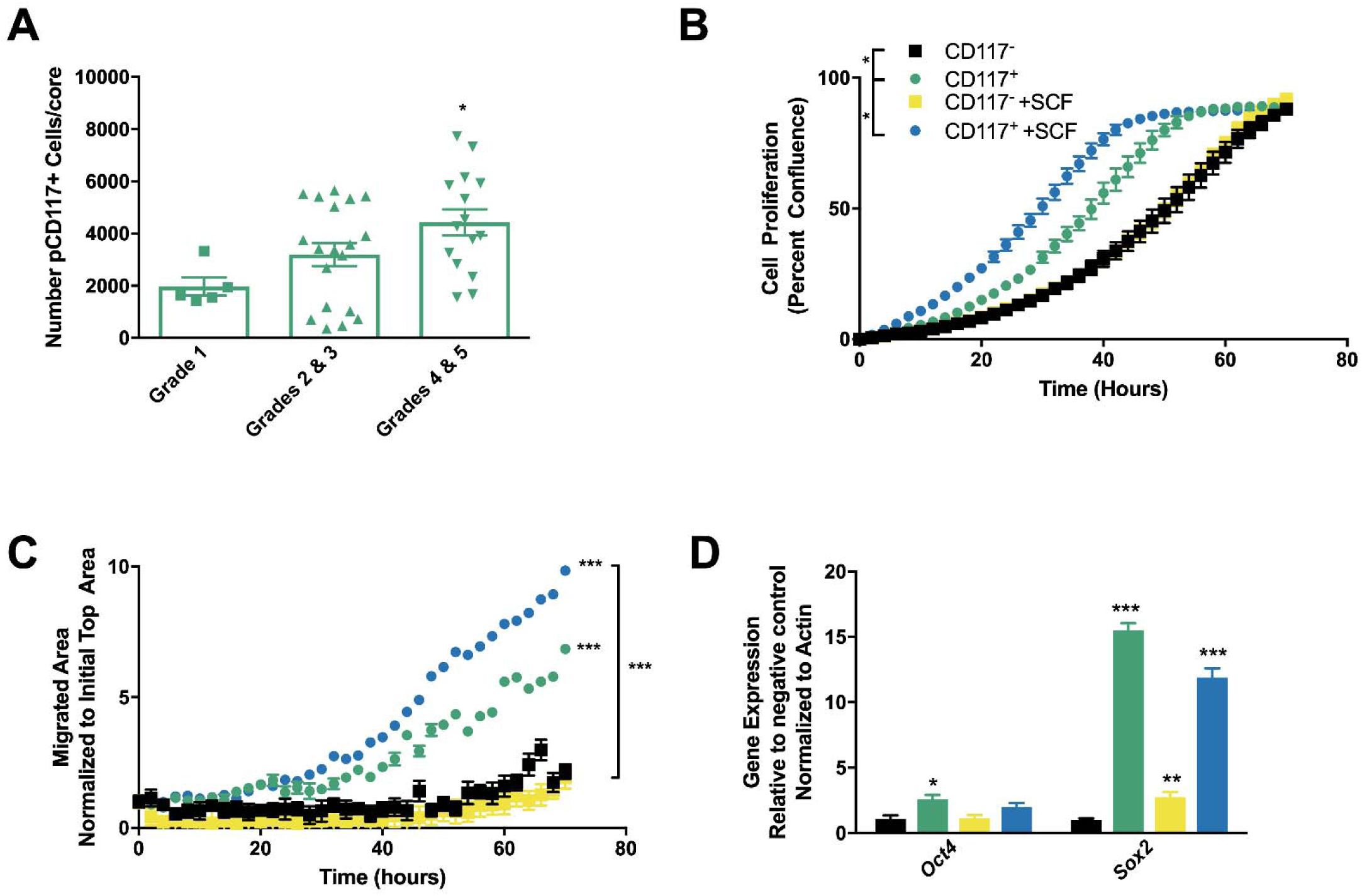
CD117 Activation by SCF Stimulates Aggressiveness. (A) Tumor microarrays from prostate cancer patients were stained for phosphorylated (p) CD117. The number of positive cells per core were counted as mean ±SEM (*n*=5-20) and separated by Gleason score. (B-D) Sorted LNCaP-C4-2 cells were treated with 50 ng/mL stem cell factor (SCF) to activate CD117. (B) Proliferation was assessed by live-cell imaging and represented as mean percentage confluence ±SEM (*n*=12). (C) Chemotaxis through pores towards SCF was measured by live-cell imaging and represented as mean number of cells migrated to the initial numbers seeded ±SEM (*n*=8). (D) Gene expression of *Oct4* and *Sox2* were measured after SCF treatment and represented as mean foldchange ±SEM (*n*=4). * represents p<0.05, ** represents p<0.01, and *** represents p<0.005 by one-way ANOVA.

Next, we examined how CD117 activation by SCF alters prostate cancer cell aggressiveness. First, we treated our sorted CD117+ and negative cells with SCF and found that SCF increased proliferation of CD117+ cells by live-cell imaging (1.4-fold at hour 40) but had no significant effect on the negative populations (Figure 5B). We then tested how CD117+ cell activation by SCF would change invasion. Sorted cells were placed in a transwell system with SCF in the bottom chamber (Figure 5C). CD117+ cell invasion was 1.4-fold higher towards SCF than media alone at experimental termination. Thus, SCF functions as a chemoattractant factor for CD117+ cells inducing migration. Finally, SCF treatment effects on CSC gene expression was examined. CD117+ cells have much higher CSC gene expression compared with the negative population. SCF treatment stimulated small, but insignificant decreases in *Oct4* and *Sox2* expression in CD117+ cells and an increase in *Sox2* expression only in negative cells (Figure 5D). Taken together, these data demonstrate that CD117 activation induces prostate cancer progression as shown by proliferation and migration.

### CD117 Inhibition Prevents Prostate Cancer Progression

Since CD117 activation induced prostate cancer progression and migration, we next examined the effects of treatment with the tyrosine kinase inhibitors (TKIs): sunitinib, imatinib, and ISCK03. Sunitinib targets several pathways and receptors including PDGFR and VEGFR in addition to CD117.[21],[43]–[45] Imatinib targets CD117, in addition to BCR-Abl, RET, and PDGFR.[43],[46] ISCK03 is a cell-permeable CD117 specific inhibitor that blocks SCF-induced phosphorylation.[47]–[49] In proliferation studies, sunitinib had the greatest reduction of proliferation for CD117+ cells (85% at 70 hours), while ISCK03 treatment resulted in a 50% reduction in proliferation and imatinib treatment did not affect proliferation (Figure 6A). We next examined the effect of TKIs on growth in 3D. In contrast to proliferation in monolayer, imatinib had the strongest effect inhibiting sphere growth 66%, while sunitinib induced 50% inhibition, and ISCK03 had minimal effect at hour 136 (Figure 6B). Taken together, these data demonstrate that TKIs decrease CD117 induced proliferation and sphere formation, but the effects of the individual TKIs are dependent on the culture conditions. Overall our data demonstrate that CD117 activation plays a key role in prostate cancer, progression, migration, and resistance to TKIs.

**Figure 6.**
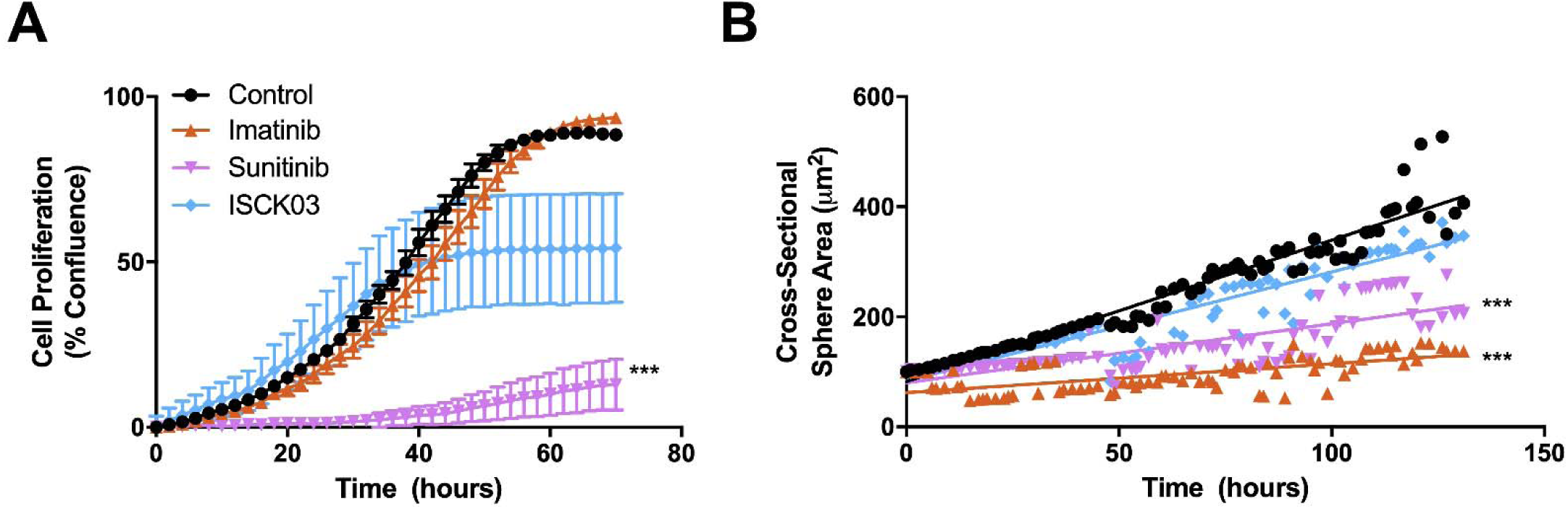
Inhibition of CD117 Decreased Proliferation and Sphere Formation. Sorted LNCaP-C4-2 cells were treated with the broad tyrosine kinase inhibitors imatinib and sunitinib, as well as, ISCK03, a CD117-specific inhibitor. (A) Proliferation was measured by live-cell imaging and represented as mean percent confluence ±SEM (*n*=6). (A) Sphere formation was tracked using live-cell imaging and represented as mean sphere area (*n*=12). *** represents p<0.005 by one-way ANOVA.

## DISCUSSION

In this study, we determined how the CTC marker CD117 expression and activation affected prostate cancer progression. CD117 expression on human prostate cancer cells induced increased proliferation, migration, and sphere formation. CD117+ cells expressed stemness marker genes and generated tumors in serial tumor initiation studies indicating that CD117-expressing cells represent a CSC. Beyond expression, CD117 activation was associated with increased cancer severity. SCF activation of CD117 stimulated further proliferation and invasion but had no additional effect on CSC genes. Conversely, CD117 inhibition by TKIs diminished cell proliferation and sphere formation. Our data suggest that CD117 activation drives prostate cancer progression, invasion, and TKI resistance through its induction of the CSC phenotype.

We demonstrate that CD117 expression induces prostate cancer progression, and its activation increases with cancer severity. CD117 expression and overactivation is found in several cancers including gastrointestinal stromal tumors (GIST), acute myeloid leukemia, and melanoma.[21],[24],[50]–[52] CD117 is best studied in GIST, which carries CD117 activating mutations. In GIST, CD117 expression and activation is associated with worse prognosis and bone metastasis.[53]–[56] However, activating mutations have only been found in GIST despite increased expression in prostate and ovarian cancers among others.[21],[52],[57] CD117 expression in many cancers is associated with shorter disease survival and metastasis and is increased with cancer progression in prostate cancer patients with the highest levels of CD117 staining seen in bone metastases.[21],[57]–[59] However, the opposite is true in myeloid/erythroid cancers, whose patients have improved or unchanged prognosis when CD117 is expressed.[60] This may be partially due to biological differences between hematologic malignancies and solid tumors in which some genes and miRNA have opposite functions.[61] Further, in solid tumors, cancer cells are often the result of dedifferentiation whereas in hematological malignancies cancer cells are often transformed HSCs or from earlier lineage cells. Thus, the differences in CD117 function in these tumors may relate to the degree of “stemness.” Our data indicate that the CD117 subpopulation represents a prostate CSC. This finding is supported by data demonstrating that CD117 expressing cells in the normal prostate are also stem-like. CD117+ prostate stem cells were found in all murine prostate lobes and both the luminal and basal compartments. Prostates generated from a single CD117+ cell contained neuroendocrine synaptophysin-positive cells and expressed probasin and Nkx3.1.[22] CD117 is a well-known stem-cell marker for normal hematopoietic cells, and its expression declines as cells lose their plasticity during differentiation.[62]–[64] Further, in other cancer types, including lung and ovarian cancers, CD117 expressing cells exhibited CSC characteristics including self-renewal.[65]–[67] In osteosarcomas, CD117 is expressed on CSCs and confers resistance to chemotherapies.[68] In that study, CD117+ osteosarcoma cells formed spheres and successfully initiated tumors after serial transplant similar to our findings with the prostate cancer cell line. Further, in accordance with previous studies for prostate cancer, CD117 expression was higher in metastatic osteosarcoma tumors. Thus, CD117 expression appears to correlate with an ability to metastasize to bone. Our prior study demonstrating CD117 expression on CTCs, in combination with this study demonstrating that CD117+ cells are tumor-initiating cells, suggests that CD117 likely plays an important role in driving bone metastasis although future studies are needed to examine a causal link.

CD117+ cell escape from the primary tumor could be caused by chemotaxis towards SCF as demonstrated by our *in vitro* data. SCF plays an important role in the homing to and maintenance of HSCs in the bone microenvironment.[58],[64],[69],[70] Bone marrow niche cells secreting SCF include perivascular cells, endothelial cells, pericytes, mesenchymal stem cells, megakaryocytes, and stromal cells.[71],[72] Additionally, osteoblasts produce SCF and control CD117 expressing HSC numbers near trabeculae.[73] Correspondingly, SCF deletion in endothelial cells or pericytes leads to HSC depletion in bone marrow.[71],[74],[75] SCF also plays an important role in cancer and metastasis. Bone marrow stromal cells and prostate cancer express membrane SCF and release the soluble form. However, bone marrow stromal cells express much higher levels of the soluble SCF.[58] These data indicated that SCF in the bone marrow might function as a chemoattractant stimulating prostate cancer bone metastasis as already demonstrated during Ewing’s sarcoma metastasis to bone.[76] Interestingly, LNCaP-C4-2 prostate cancer cells contain a subpopulation of CD117+ cells, while the LNCaP parental subline does not. It is likely that the exposure of the LNCaP cells to bone marrow during the generation of the C4-2 subline induced CD117 expression.[77] This was shown with the PC3 cells that once exposed to bone marrow cells started to express CD117.[58] In other cell types, SCF treatment can induce CD117 expression but can also induce internalization of the receptor.[41],[78],[79] These data may explain why SCF treatment stimulated increased *Sox2* expression in the negative population and had little effect on the CD117+ cell population gene expression. Further studies are needed to examine how SCF treatment might promote CD117 expression in negative cells. Binding of SCF to the CD117+ cells may confer therapeutic resistance due to internalization of the receptor or via competition with TKIs, such as imatinib, for the CD117 activation site.[80]

Our data demonstrate that CD117+ prostate CSCs do not have a strong response to TKI treatment and that their response is context dependent. The TKIs imatinib and sunitinib were developed for and have a higher specificity for other tyrosine kinases. The lack of CD117+ cell response may be one reason for the failures of these TKIs in clinical studies.[81] Further, for prostate cancer, TKIs were given to patients with castration-resistant metastatic disease, which may be too late for the treatments to show efficacy. However, cabozantinib treatment in CRPC patients demonstrated tumor reduction and smaller bone metastases,[82] which was not tested in this study, had a higher specificity for CD117 than either imatinib or sunitinib.[21] CD117 expressing osteosarcoma cells are resistant to doxorubicin.[68] CD117 mutations in GIST are responsible for resistance to TKI treatment. 14% of GIST patients are initially resistant to imatinib, and 50% develop resistance within two years of treatment. For most patients, sunitinib will then be used and effective unless one mutation, D816H/V, is present which is resistant to both TKIs. Imatinib works better on inactive CD117 and prevents activation, but does not bind to activated CD117.[83] Further, SCF in the bone microenvironment alters the efficacy of treatments, especially TKIs, which have off-target effects in bone.[84] TKIs induce osteonecrosis in the jaw even when combined with bisphosphonates.[85] One possible reason may be stimulation of osteoclast and osteoblast differentiation by SCF. Another confounding factor is that tumor microenvironment SCF induces imatinib resistance by competing for the binding site with a higher affinity for CD117.[80] Thus, the high levels of SCF in the bone microenvironment may prevent TKI treatment from working properly on CD117 expressing prostate cancer cells, the numbers of which increase after exposure to the bone marrow.[58] The failures of prior TKI research may be due to the testing of inhibitors in patients who had already developed bone metastases.[81] This may be too late to alter progression and TKIs could be more effective earlier in the metastatic process. Further, failures may be due to the low specificity of current TKIs for CD117.[21]

In combination with our prior data showing a CD117+ CTC population in advanced prostate cancer patients, the data from this study demonstrate that CD117 expression in primary tumors and on CTCs may distinguish advanced cancer patients likely to have more aggressive tumors leading to recurrence and metastasis. Further research is required to verify that CD117 expressing cells represent CSCs in patients and that these cells initiate bone metastases, which would require the development of new bone metastases models.[86] In summary, we demonstrate that the CD117 subpopulation of LNCaP-C4-2 prostate cancer cells represents CSCs driving progression, migration, and TKI resistance.

## AUTHOR CONTRIBUTIONS

Investigation, Writing, Review, and Editing: BMF, BAK, CDL, CJS, KSH, KW, LS, MEM, PLE; Formal analysis and Visualization: BMF, BAK, CDL, KSH, LS, MEM; Resources: CDL, KW; Conceptualization, Funding acquisition, Project administration, Supervision: BAK

## ACKNOWLEDGMENTS

We thank Brandi Bickford from the Virtual Microscopy Core for her assistance with slide scanning, Jolyn Turner from the Cell Engineering Shared Resource for assistance generating the cell lines, and Dr. Taylor Peak for assistance with TMA staining. We thank Dr. Skip Heston, Dr. Magda Grabowska, Dr. Simon Hayward, and Dr. Leland Chung for the gift of the cell lines.

